# Comparative analysis of cellular immune responses in conventional and SPF Baboons (*Papio spp.)*

**DOI:** 10.1101/455881

**Authors:** Elizabeth R Magden, Bharti P. Nehete, Sriram Chitta, Lawrence E. Williams, Joe H Simmons, Christian R. Abee, Pramod N. Nehete

## Abstract

Baboons (*papio spp.*) have served as a successful model of human disease such as cardiac and respiratory, infectious, diabetes, genetics, immunology, aging, and xenotransplantation. The development of an immunologically defined specific-pathogen free (SPF) baboon model has further advanced research, especially with studies involving the immune system and immunosuppression. In this study, we compare normal immunological changes of peripheral blood mononuclear cell (PBMC) subsets, and their function in age-matched conventional and SPF baboons. Our results demonstrate that both groups have comparable numbers of different lymphocyte subsets, but there are phenotypic differences in central and effector memory T cells subsets that are more pronounced in the CD4+ T cells. Despite equal proportions of CD3+ T cells among the conventional and SPF baboon groups, PBMC show higher proliferative responses to mitogens PHA and PWM and higher IFN-γ producing cells to Con A and PWM in the conventional group. Plasma levels of the inflammatory cytokine TNF-α were significantly higher in SPF baboons. Exposure of PBMC from conventional baboons to various Toll like ligands (TLR ligands) TLR-3, TLR-4 and TLR-8 show higher IFN-γ producing cells while PBMC from SPF baboons stimulated with TLR-5 and TLR-6 ligand show higher IFN-γ producing cells. These findings suggest that while the lymphocyte subsets in conventional and SPF baboons share many phenotypic and functional similarities, specific differences exist in immune function of lymphocytes which could impact the quality and quantity of innate and adaptive immune responses. These differences should be considered for better experimental outcomes, specifically in studies measuring immunological endpoints.

## Introduction

As a disease model, baboons have a lot to offer. In comparison to other nonhuman primates, baboons are large. Their large size allows researchers to study organ transplantation, medical devices, and collect ample fluid and tissue samples. Baboons also share approximately 96% genetic homology with humans, making them a more relevant animal model than the typical laboratory rodent [1]. Baboon research has focused on cardiac disease (coronary heart disease, hypertension, and atherosclerosis), respiratory diseases (*Bordetella pertussis* and respiratory syncytial virus), xenotransplantation, reproductive and neonatal physiology, diabetes, genetics, infectious disease, immunology and vaccine development [2-9]. Baboons are an excellent model for vaccine development as their immune system shares similarities with the human immune system since baboons have the same immunoglobulin G (IgG) subclasses 1, 2, 3, and 4 as humans [10].

Most baboons are maintained under conventional conditions (non-SPF). Under these conventional conditions, baboons harbor a number of adventitious virus infections that typically do not cause disease in immune competent animals and are often considered part of the animal’s normal flora including a complete complement of herpesviruses and retroviruses. Baboons in specific pathogen-free (SPF) colonies have been bred to eliminate many of these agents. The bioexclusion list for SPF animals can vary by institution. The SPF baboons examined in this study are negative for 21 agents including multiple viruses (all known herpesviruses, retroviruses, polyomaviruses, paramyxoviruses, and orthopoxviruses), internal parasites (*trichuris* spp., *stronglyoides spp*., and *giardia* spp.), and two bacteria species (*Mycobacterium* tuberculosis complex bacteria and *Bordetella* spp.). Retroviruses and herpesviruses are of particular concern to biomedical research due to their ability to influence the immune system and to persist in the host after the initial infection event. Because monkey herpesviruses are closely related to their human virus orthologues, testing of human herpesvirus vaccines in conventional nonhuman primates is problematic due to antigenic cross reactivity between human and enzootic simian herpesviruses. In addition, the ability of most baboon herpesviruses to infect human cells *in vitro* raises the risk of zoonotic infection [11].

These risks highlight the need for SPF baboons in immunological studies and vaccine development. Studies thus far have primarily utilized conventional baboons, which raises the question of if there is a difference in cellular immune responses between SPF and conventional baboons. In this study, we compared normal immune cell parameters between age-matched conventional (non-SPF) and SPF baboons.

## Materials and methods

### Ethics Statement

This research was conducted at the AAALAC (Association for Assessment and Accreditation of Laboratory Animal Care) accredited Michale E. Keeling Center for Comparative Medicine and Research (Keeling Center), at the UT MD Anderson Cancer Center, Bastrop, TX (UTMDACC). All blood samples were collected as part of routine veterinary medical examination and according to the provisions of the Animal Welfare Act, PHS Animal Welfare Policy, and the principles of the NIH Guide for the Care and Use of Laboratory Animals. All procedures were approved by the Institutional Animal Care and Use Committee at the UT MD Anderson Cancer Center.

### Animal, Diet, and Blood collection

In the present study, we compared immune responses of conventional and SPF baboons, each cohort consisting of 25 females in the 7-13 year old age range that were socially housed in indoor/outdoor runs or Primadomes™ [12] at the UTMDACC, Bastrop, Texas. Animals have *ad libitum* access to Lab Diet^®^ #5045 high protein nonhuman primate diet and water. In addition, they are fed fresh fruits and vegetables daily, as well as regular enrichment items such as forage, seeds, peanuts, raisins, peanut butter, and frozen juice cups. Subjects are also provided with destructible enrichment manipulanda and different travel/perching materials on a rotating basis to promote the occurrence of typical species behavior. The colony management practices include a comprehensive veterinary program to assess baboon health and psychological well-being along with the daily environmental enrichment opportunities. We used well-established criteria from the literature in choosing healthy animals for our study, all were free of illness [13].

SPF baboons are defined by Wolf et al [14] as being absent of the following bioexclusion list of pathogens: Herpesviruses (Herpesvirus papio 1 and 2, Simian varicella zoster virus, Baboon cytomegalovirus, Human herpes virus 6, Baboon rhinovirus), Retroviruses (Simian foamy virus, Simian retrovirus D, Simian immunodeficiency virus, Simian T lymphotropic virus), Polyomaviruses (Simian virus 40 and SA12), Paramyxovirus (Morbillivirus measles), Orthopoxvirus (Monkeypox virus), Arterivirus (Southwest baboon virus 1), Internal Parasites (*Trichuris trichuria-*whipworms), *Stongyloides* sp. *–*threadworms*, Giardia* sp.), Blood parasites *(Babesia sp.)* and Bacteria (Mycobacterium tuberculosis spp. bacteria and *Bordetella* sp.).

### Clinical and laboratory assessment of study animals

Animals are observed twice daily by the veterinary staff as part of the comprehensive veterinary care program. Animals are sedated for biannual physical examinations and as needed to treat illness or injury. Blood samples are collected from a peripheral vein and analyzed for a complete blood count (Siemens Advia 120 Hematology Analyzer, Tarrytown, NY) and serum chemistry profile (Beckman Coulter AU680^®^ Chemistry Immuno Analyzer, Brea, CA). The absolute number of lymphocytes, obtained from hematological analysis, was used in converting the frequency of the lymphocyte population obtained from FACS analysis, in order to get the absolute number of the lymphocyte subset populations.

### Study Groups, Collection of samples, and PBMC preparation

Blood samples (10 mL) were collected via venipuncture of the femoral vein from 50 baboons (conventional and SPF, each n=25) between 9 a.m. and 11 a.m. in EDTA anticoagulant tubes after the animals were anesthetized with ketamine (10 mg/kg intramuscularly, Vedco Inc., Saint Joseph, MO). Blood samples were processed at the Keeling Center within 2-4 hours of collection. Plasma was separated by centrifugation and stored at -80°C till further use. PBMCs were isolated by Ficoll-Hypaque density gradient separation as described previously ([15, 16]. Erythrocytes were removed by osmotic lysis in ACK lysing buffer (Life Technologies, Grand Island, NY), and the remaining nucleated cells were washed twice with RPMI supplemented with 10% fetal bovine serum (FBS, Atlanta Biological, Flowery Branch, GA) and used for immune assay.

### Flow cytometry

A series of commercially available human monoclonal antibodies that cross-react with NHP mononuclear cells were used in flow cytometry analyses, as described previously [15-18]. Briefly, 100 µL of whole blood from each sample was added to each 12-mm × 75-mm polystyrene test tube (Falcon, Lincoln Park, NJ) containing a panel of monoclonal antibodies CD3 Percp (clone SP-34), CD8 PE (clone SK1), CD16 FITC (clone 3G8) and CD20 APC (clone L27) (all from BD Biosciences, San Diego, CA) and incubated for 15 min at room temperature in the dark. Red blood cells were lysed with 1× FACS lysing solution (Becton Dickinson, San Diego, CA), diluted according to the manufacturer's instructions. The samples were washed thoroughly in 1× phosphate-buffered saline (PBS) by centrifugation; cell sediments were then suspended in 1% paraformaldehyde buffer (300 µL), and cells were acquired on a 4-color flow cytometer (FACSCalibur; BD Biosciences, San Jose, CA). Lymphocytes that were gated on forward scatter versus side scatter dot plot were used to analyze CD3^+^, CD4^+^(CD3+CD8-),CD8^+^ (CD3+CD8+) T-cell and CD20^+^ B-cell lymphocyte subsets with use of FlowJo software (Tree Star, Inc., Ashland, OR).

For NK and NKT cell analysis, a separate tube of 100 µL of blood was used for staining with the combination of anti-CD3 (Percp, clone SP-34); CD8 PE (clone SK1) and anti-CD16 (FITC clone 3G8), (BD Pharmingen, San Jose, USA) antibodies, as described above. The stained cells were acquired with FACSCalibur (Becton Dickinson) and analyzed with use of FlowJo software (Tree Star, Inc., Ashland, OR, USA).

For the analyses of T cell memory subsets, 100 µL of EDTA-preserved whole blood was stained with anti-CD3 (FITC, clone SP34-2, BD Biosciences, San Jose, CA), anti-CD4 (PE, clone L200, BD Biosciences), anti-CD28 (PerCpCy5.5 BD Biosciences), and antiCD95 (APC, clone DX2, BD Biosciences) and process as mentioned above. Both compensation controls and fluorescence minus one (FMO) controls were utilized. Results were acquired on a FACSCaliber (BD Biosciences) and analysed using FlowJo software (Tree Star, Inc., Ashland, OR).

### In Vitro mitogen stimulation

PBMCs freshly prepared from whole blood collected in an EDTA tube were more than 90% viable, as determined by the trypan blue exclusion method, and for each immune assay we used 10^5^ cells/well. The proliferation of PBMCs was determined by the standard MTT dye reduction assay, as previously described [18-21]. Briefly, aliquots of PBMCs (10^5^/well) were seeded in triplicate wells of 96-well, U-bottom plates and individually stimulated for 48 h with the mitogens phytohemagglutinin (PHA), concanavalin-A (Con A), lipopolysaccharide (LPS), and pokeweed mitogen (PWM) (Sigma, St Louis, MO), each at a final concentration of 5 µg/mL. The culture medium without added mitogens served as a negative control. After culture for 48 h at 37°C in 5% CO_2_, 175 µL of medium was replaced with 15 µL of freshly prepared MTT dye (5 mg/mL in PBS). After 4 h of incubation, medium was then replaced with 100 µL of 0.04 N acidified isopropanol (Sigma). After 30 min of incubation at room temperature for color development, the plate was read at 490-nm ELISA plate reader (Victor, PerkinElmer, Shelton, CT). Results were expressed as optical density (OD) after blank (i.e., medium only) subtraction. Reported values were the mean of 3 replicates. The optimal concentration of mitogen, number of PBMCs, and incubation time were previously standardized in our laboratory from PBMCs isolated from healthy animals.

### ELISpot Assay for Detecting Antigen-specific IFN-γ-producing Cells

Isolated PBMCs were stimulated individually with the mitogens Con A, PHA and PWM (each at 2 μg/mL final concentration) to determine the numbers of IFN-γ-producing cells by the ELISpot assay using the methodology reported earlier [22, 23]. Briefly, aliquots of PBMCs (10^5^/well) were seeded in duplicate wells of 96-well plates (polyvinylidene difluoride backed plates, MAIP S 45, Millipore, Bedford, MA) pre-coated with the primary antibody to IFN-γ and incubated with PHA, Con A, and PWM. After incubation for 24 hr at 37°C, the cells were removed and the wells were thoroughly washed with PBS and developed as per protocol provided by manufacturer. Purple colored spots representing individual cells secreting IFN-γ were counted by an independent agency (Zellnet Consulting, New Jersey, NJ) using the KS-ELISPOT automatic system (Carl Zeiss, Inc. Thornwood, NY) and the data shown as number of IFN-γ spot forming cells (SFC) for 10^5^ input PBMCs. Responses were considered positive when the numbers of spot forming cells (SFC) with the test antigen were at least five SFC and also were five SFC above the background control values from cells cultured in the medium alone.

### Cytokine multiplex assays

The concentration of cytokines IFN-γ, interleukin 2 [IL-2], IL-6, IL-10, IL-12 (p40), and tumor necrosis factor α (TNF-α), in plasma were measured with use of a NHP Multiplex Cytokine Kit (Millipore Corporation, Billerica, MA), as described previously [18]. Briefly, EDTA-preserved plasma samples were centrifuged (1200 × *g* for 10 min), and aliquots were frozen at −80°C until used. On the day of assay, plasma samples were thawed and pre-cleared by centrifuging at 1200 × *g* for 5 min. The 96-well plates provided in the kit were blocked with assay buffer for 10 min at room temperature and washed, and 25 µL of standard or control samples were added in appropriate wells. After 25 µL of beads were added to each well, the plate was incubated on a shaker overnight at 4°C. The next day, after the plate was washed twice with wash buffer, it was incubated with detection antibody for 1 h, followed by another incubation with 25 µL of streptavidin-phycoerythrin for 30 min. All incubation and washing steps were performed on a shaker at room temperature. After the plate was washed twice with wash buffer, 150 µL of sheath fluid was added to each well, and cytokines were measured by acquiring beads on the BioPlex 200 system (Luminex X MAP technology). Fluorescence data were analyzed with use of Bio-Plex Manager 5.0 software (Bio-Rad, Hercules, CA). The minimum detectable concentration was calculated by the Multiplex Analyst immunoassay analysis software from Millipore. The minimum detectable concentrations in pg/mL for the various cytokines were as follows: IFN-γ (2.2), IL-2 (0.7), IL-6 (0.3), IL-10 (6.2), IL-12(p40) (1.2), and TNF-α (2.1).

### Ex vivo induction of cytokines by TLR ligands

PBMCs obtained after centrifugation of blood using a density gradient were washed with PBS. Aliquots of 1 × 10^5^ cells were re-suspended in culture medium RPMI-1640 (Hyclone Laboratories) were dispensed in each well of a 96-well plate. The culture medium used was free of detectable endotoxin (<0.1 EU/mL) and all other solutions were prepared using pyrogen-free water and sterile polypropylene plastic ware. The cells were then incubated with or without TLR ligands (TLR-1 Pam3CSK4, TLR-3 Poly I:C, TLR-4 LPS, TLR-5 Flagellin, TLR-6 FSL1, RLR-8 ssRNA40/LyoVec.) all from; Invivogen (Invivogen Corp, San Diego, CA, USA) at 1 μg/mL for 24 hr. at 37°C in a 5% CO_2_ atmosphere. Cells were used for IFN-γ Elispot as described above.

### Statistical analysis

For statistical analysis, samples were grouped according to conventional and SPF animals and unpaired two-tailed *t* test analyses were performed for the data. Results, expressed as mean ± standard deviation and statistic significances, were shown in the corresponding figure and figure legends. An F test for equal variances was done to ensure that the groups had equal variances before the *t* tests were run. The *p* values <0.05 were considered statistically significant. All statistical analyses were conducted using GraphPad Prism^®^ 6.00 (GraphPad Software, San Diego, California USA).

## Results

Conventional baboons harbor a number of adventitious infectious agents common to nonhuman primate species, including many of the pathogens on the SPF bioexclusion list noted above. Since these agents have been eliminated from the SPF baboon colony, we assessed if their absence would result in changes to the complete blood count (CBC) or blood chemistry. We examined the CBC and blood chemistry, and found no significant differences between conventional and SPF baboons as has been previously reported [14].

### Expression of major lymphocyte subsets in peripheral blood of conventional and SPF baboons

Peripheral blood samples collected from the conventional and SPF groups of baboons were analyzed by flow-cytometry to enumerate different lymphocyte subsets. A gating strategy for phenotypic analysis of T and B cells is shown in Fig 1A. We observed no significant differences in the absolute numbers of CD3^+^ (T cells), CD4+ (T helper cells), CD8+ (T suppressor cells), CD4+CD8+ (double positive T cells), and CD20^+^ (B cells) between conventional and SPF baboons (Fig 1B).

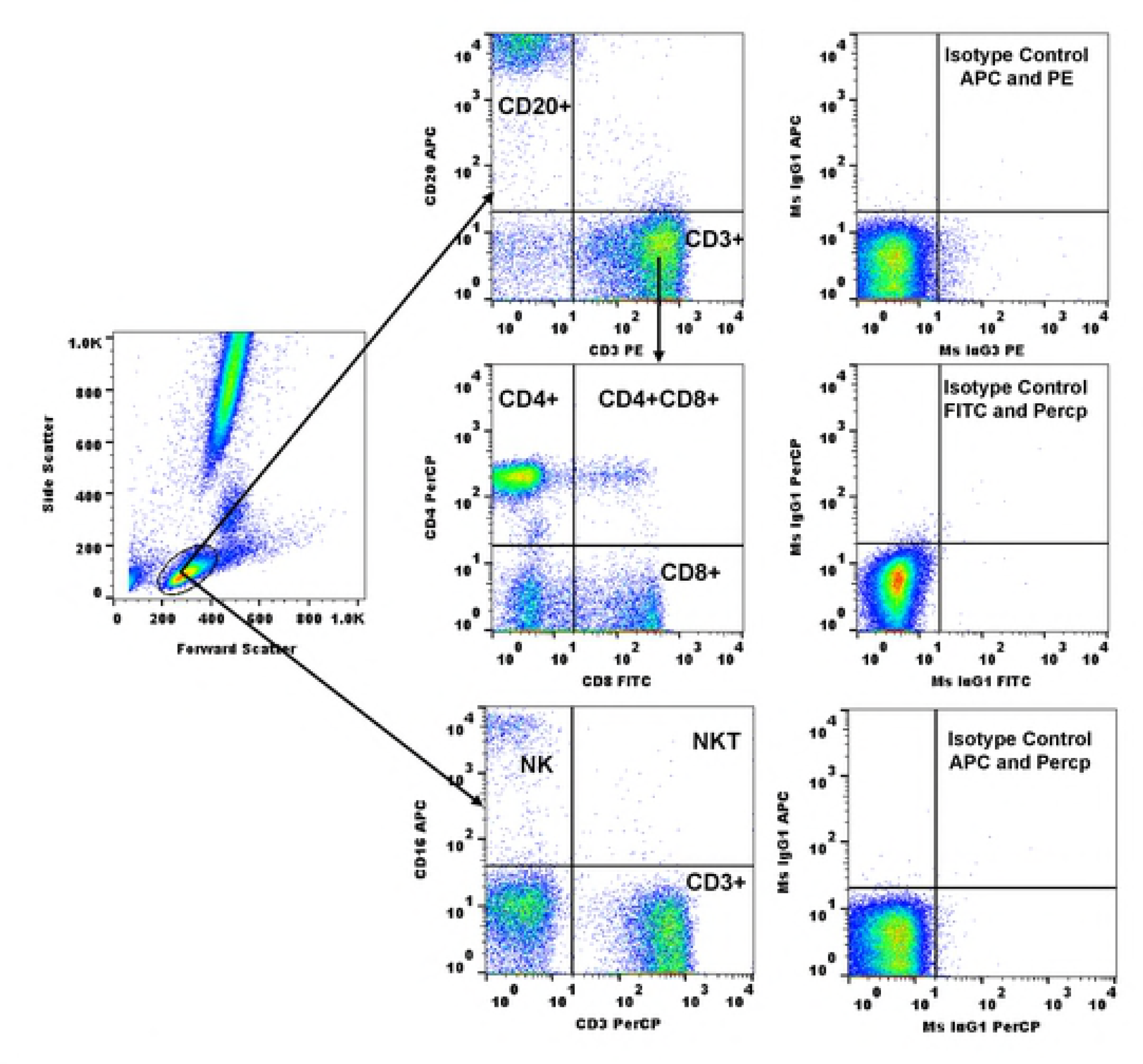
**Fig 1 (A): Gating scheme for phenotype analyses of the various cell markers in the peripheral blood from a representative animal** The lymphocytes and monocytes were first gated based on forward scatter (FCS) versus side scatter (SSC), and then CD3^+^ T cells, CD4^+^, CD8^+^,CD4+CD8+, CD20^+^ B cells and CD16+ NK and NKT cells were positively identified. The specificity of staining for the various markers was ascertained according to the isotype control antibody staining used for each pair of combination markers, as shown. **Fig 1 B and 1C: Phenotype analysis of lymphocytes in conventional and SPF baboons** Aliquots of EDTA whole blood were stained with fluorescence-labeled antibodies to the CD3^+^, CD4^+^, CD8^+^, CD16+ and CD20^+^lymphocytes and analyzed for T-cell subpopulations in conventional and SPF baboons. Analysis of lymphocytes, CD3+ and its subsets and CD20 are shown in figure 1B and CD16+ and its subsets are shown in figure 1C. Values on the Y-axis are absolute lymphocytes cells. P values were considered statistically significant at \**p*<0.05.

Expression of NK and NKT subsets in peripheral blood of conventional and SPF baboons is shown in Fig 1C. NK cells are an important component of the innate immune response that has fundamental roles in the defense against certain cytopathic viruses, primarily herpes viruses [24]. Baboon NK cells are identified as CD3^-^CD16^+^ cells and subdivided based on the expression of CD8. We observed no significant difference in NK and NKT cells between conventional and SPF baboons. However, CD8+NKT cells were significantly higher in SPF than conventional baboons (t=2.09, df=33, *p*<0.05, η2=0.12) (Fig 1C).

### Phenotype analysis of memory markers in conventional and SPF baboons

To investigate naive and memory T-cell subsets in conventional and SPF baboons, we used surface expression of CD95 and CD28, most consistently used in both human and rhesus monkeys [25, 26]. The different memory and naïve subsets of CD4+ and CD8+ T cells were enumerated by flow cytometry using the gating strategy shown in Fig 2A. The naïve (Tn), effector (TEM), and memory (TCM) subsets of CD4+ and CD8+ T cells were also enumerated by using co-stimulatory (CD28) and pro-apoptotic antigen (CD95) markers of expression using the gating strategy, as reported previously [26]. Naive T cells were identified by intermediate to high expression of CD28 and a lack of CD95; memory phenotype T cells acquire surface expression of CD95 and can further be divided into CD95^+^CD28^+^ central-memory cells and CD95^+^CD28^–^ effector-memory cells, hypothesized to be terminally differentiated. Among the conventional and SPF baboon groups of animals, within the CD4+ T cells subsets, significantly higher numbers of CD4+ central and effector memory T cells were observed in conventional baboons relative to those in the SPF groups (t=2.09, df=33, *p*<0.05, η2=0.12). In CD8+T cell subsets, significant higher numbers of CD8+ effector memory T cells were observed in conventional baboons compared to SPF baboons (t=2.09, df=33, *p*<0.05, η2=0.12) (Fig 2B). However, as shown in Fig 2B, no significant differences were observed for the naïve CD4+ and CD8+ T cells and also central memory subsets of CD8+ T cells (Fig 2B).

**Fig 2 (A): Gating scheme for Naïve and memory markers in the peripheral blood from a representative animal**

The lymphocytes were first gated based on forward scatter (FCS) versus side scatter (SSC). The T cells were then positively identified by CD3 expression followed by the detection of the CD4+ CD8-(CD4+ T cells) and CD4-CD8+ (CD8+ T cells) populations within the CD3+ T cells. On the basis of CD28 and CD95 expression, the CD4+ and CD8+ T cells were further differentiated into naive (Tn CD28+ CD95-), central memory (Tcm CD28+ CD95+) and effector memory (Tem CD28-CD95+) subsets. The specificity of staining for the different markers is ascertained based on fluorescence minus one (FMO) controls shown and as described in the methods section.

**Fig 2 (B): Analyses of memory T cell subpopulations**

Blood samples from conventional and SPF baboons were stained and analysed for T cell subpopulations by flow cytometry as described in the methods section. Absolute numbers of naïve (CD28+ CD95-), central memory (CD28+CD95+), and effector memory (CD28-CD95+) subsets of CD4 were compared between the conventional and SPF baboons groups. The results shown are an average of 10 baboons in each group and *P* < 0.05 was considered statistically significant.

### Proliferative responses

The proliferation of PBMC samples from the conventional and SPF baboons were measured as the reduction of tetrazolium salts using MTT dye and O.D. was expressed as % viability (Fig 3). Absorbance values that are lower than the control cells indicate a reduction in the rate of cell proliferation. Conversely, a higher absorbance rate indicates an increase in cell proliferation. The proliferative responses to PHA (t=3.55, df=38, *p*<0.0001, η^2^=0.24), and PWM (t=2.91, df=38, *p*<0.001 η^2^=0.19) were significantly higher in conventional when compared to those in the SPF baboons (Fig 3), but a significantly higher response was observed with Con A (t=9.11, df=38, *p*<0.05, η^2^=0.69) in SPF baboons compared to conventional baboons (Fig 3).

**Fig 3: Proliferative response of PBMC to mitogens in conventional and SPF baboons**

We used PBMCs that were isolated from blood samples of baboons to determine the proliferative response to various mitogens, using the standard MTT dye reduction assay. Proliferation responses were measured as optical density (OD) and expressed as % viability over blank (i.e., medium only). P values of <0.05 were considered statistically significant.

### ELISPOT assay for detecting mitogen-specific IFN-γ producing cells in conventional and SPF baboons

PBMCs were analyzed for the numbers of cells producing IFN-γ in response to stimulation with PHA, PWM, and LPS by the cytokine ELISPOT assay. As shown in Fig 4, conventional baboons showed significantly higher numbers of IFN-γ–producing cells in response to stimulation with Con A (t=3.88,df=42,p<0.05,η^2^=0.26), PWM (t=4.23,df=38,p<0.05, η^2^=0.32) and LPS (t=4.14,df=42,p<0.05, η^2^=0.29). Although no statistically significant differences were observed for the numbers of IFN-γ–producing cells in response to stimulation with PHA between the two groups, conventional baboons had a greater response in comparison to SPF baboons (Fig 4).

**Fig 4: IFN-γ Elispot response to mitogens in conventional and SPF baboons**

Duplicate wells of the 96-well microtiter plates, pre-coated with IFN-γ antibody, were seeded with 10^5^ PBMC from the baboons and incubated with 5µg of each of the mitogens for 36 h at 37°C. At the end of the incubation period, the wells were washed and stained with biotynylated second IFN-γ antibody. The total number of spot forming cells in each of the mitogen-stimulated wells was counted and adjusted to control medium as background. See methods section for experiment details. P values were obtained from the student’s *t*-test.

### Circulating cytokines peripheral blood plasma

We analyzed plasma samples from conventional and SPF baboons for 6 different Th-1 (IFN-γ, IL-2, TNF-α) and Th-2 (IL-6, and IL-10) cytokines and IL-12(p40), using the bead array kit, and observed significantly higher levels of TNF-α (t=2.89, df=45, p<0.05, η^2^=0.16) in SPF compared to conventional baboons. However, other cytokines show no differences between SPF and conventional baboons (Fig 5).

**Fig 5: Cytokine Bead array (CBA) in plasma of conventional and SPF baboons**

Plasma cytokines from conventional and SPF baboons were measured using Bio-Rad 200 Luminex technology. In duplicate wells of the 96-well filter plate 25µl of plasma was incubated with 25µL of cytokine coupled beads for overnight at 4°C followed by washing and staining with biotynylated detection antibody. Results are expressed as pg/mL concentration. The minimum detectable concentrations in pg/mL for IFN-γ (2.2),TNF-α (2.1), IL-2 (0.7), IL-6 (0.3), IL-10 (6.2), IL-12(P40) (1.2), are used for considering positive responses. See methods section for experimental details. The standard deviation values did not exceed 15% of the mean value. P values were considered at p<0.05.

### Ex vivo stimulation of cytokines by TLR ligands

PBMCs (1 × 10^5^) incubated with ultra-purified TLR ligands (TLR-1,-4, -3, -5, -6- and 8) all from; Invivogen (Invivogen Corp, San Diego, CA, USA) at 1 μg/mL for 24 hr at 37°C in a 5% CO_2_ atmosphere. The cells were used for IFN-γ Elispot assay as described above and data presented as IFN-γ producing cells (Fig 6). We observed a significant increase in stimulation with TLR ligands 5, and 6 and significant decrease with TLR-4, (t=2.87, df47, p<0.0002, η^2^=0.54) and TLR-3 and TLR-8 ligands (t=2.87, df=7, p<0.05, η^2^=0.54) in baboons. (Fig 6).

**Fig 6: *Ex vivo* induction of IFN-γ production by TLR ligand**

PBMCs isolated from male conventional and SPF baboons were stimulated with TLR ligands for 24 hr and cells were measured for IFN-γ Elispot assay as described above. See method section for experiment details. P values were obtained from the student’s *t*-test.

## Discussion

The high level of genetic homology between baboons and humans along with their outbred nature and large size has made them a valuable animal model. While their research contributions are vast, we have focused on their immune system, which shares many similarities to our own immune system. We have specifically focused on the differences in immune systems between conventional and SPF baboons. Differences in immunity are likely to exist between conventional baboons with their own array of endogenous viruses, parasites, and bacteria, and SPF baboons who lack many of the pathogens found in conventional baboons. The preservation of a functional and diverse T cell population is a dynamic process controlled by exposure to antigen and cytokines.

An analysis of baboon T cells has demonstrated that CD4 and CD8 T cells can be subdivided into naïve, central memory (CM) and effector memory (EM) as reported in rhesus macaques using CD28 and CD95 as the primary cell surface markers [18, 27]. We found that absolute numbers of helper and cytotoxic T cells are the same in both conventional and SPF baboons. Our analysis of memory markers reveals that expression of CD4+ TEM, CD4+ TCM and CD8+ TEM are significantly higher in conventional versus SPF baboons. This finding parallels those observed in infant, adolescent, and adult macaques in conventional versus SPF colonies [28, 29]. In these studies, the observed differences in specific lymphocyte subsets likely contributed to the distinct cytokine responses after mitogenic T cell stimulation. Researchers noted that the conventional adolescent macaques produced higher levels of inflammatory cytokines compared to their age-matched SPF adolescent macaques.

Natural killer (NK) cells are critical to the innate immune system as the first line of defense against many virus-infected cells and tumor cells. NK cells are also important mediators of transplantation rejection reactions, as seen during baboon xenotransplantation studies [30-32]. In this study, we phenotyped baboon NK cells. Baboon NK cells are identified as CD3-CD16^+^ cells. In humans, NK cells are CD3^-^CD56^+^ (85%–90%) and can be subdivided based on the expression of CD16 [33]. One major difference between human and NHP NK cells is that baboon and macaque NK cells express high levels of CD8α, but this is not observed in human NK cells [33, 34].

To determine if the phenotypic changes we observed in this study were indicative of an altered T cell function, we measured mitogen-specific T cell responses in the plasma of conventional and SPF baboons. We found a significant increase in TNF-α in circulating cytokines of SPF baboons, while IL-2, IL-6, IL-10, IL-12(p40) and IFN-γ showed no significant differences in conventional compared to SPF baboons. The identification of noticeable differences between conventional and SPF baboons suggests that chronic infections modulate host immune development as reported in SPF and conventional rhesus macaques [35].

While the conventional and SPF baboons share many phenotypic and functional similarities of lymphocyte subsets, the findings of this study suggest that specific differences exist in lymphocyte immune function which could impact the innate and adaptive immune responses. As we plan immunological studies going forward, these differences should be considered as they could potentially impact results and outcomes for studies examining various immunologic endpoints.

## Conclusion

Baboons are invaluable models to advance our understanding of disease pathogenesis, immunity, and vaccine development due to their genetic similarity to humans and natural susceptibility to a wide variety of human pathogens. Comparative studies of the immune system of conventional and SPF baboons will enhance our knowledge of the baboon model and accelerate research on the important mechanisms associated with inflammation. This research will help to identify and validate biomarkers and develop novel vaccine strategies.

